## Supplemental Materials for "Parenting deficits in *Magel2*-null mice predicted from systematic investigation of imprinted gene expression in galanin neurons of the hypothalamus"

**Supplemental Material**

M. J. Higgs^1^, A. E. Webberley^1^, R. M. John^2^, and A. R. Isles^1*^

1 Behavioural Genetics Group, MRC Centre for Neuropsychiatric, Genetics and Genomics, Neuroscience and Mental Health Research Institute, Cardiff University, Cardiff, UK

2 School of Biosciences, Cardiff University, Cardiff, UK

* Corresponding author: A. R. Isles, Behavioural Genetics Group, MRC Centre for Neuropsychiatric Genetics and Genomics, Neuroscience and Mental Health Research Institute, Cardiff University, Cardiff CF24 4HQ, UK.

Table of Contents

Supplemental Methods	3

Specific processing steps for each unique dataset	3

Supplemental Figures	4

Supplemental Figure S1.	4

Supplemental Figure S2.	5

Supplemental Figure S3.	7

Supplemental Figure S4.	8

Supplemental Tables	9

Supplemental Table S1	9

Supplemental Table S2 A & B	10

Supplemental Table S3	11

Supplemental Table S4 A & B	11

Supplemental Table S5	12

Supplemental Table S6	12

Supplemental Table S7	13

Supplemental Table S8	13

Supplemental Table S9	13

References for Supplemental Material	14

**Supplemental Methods**

**Preoptic Area (POA) (Moffitt et al., 2018)**

The Preoptic area is of particular interest to parenting behaviour and feeding behaviour containing neural populations that contribute to a number of innate hypothalamic behaviour outputs. Moffitt et al. (2018) conducted a molecular, spatial, and functional characterisation of the murine POA and as part of this investigation they profiled 31,299 cells from male and female adult mouse POA. Alongside 21 major cell classes, 18,553 neurons were clustered into 66 distinct neuronal populations (of which 56 originated from the POA). Barcodes, Features and Matrix files were acquired from Gene Expression Omnibus through accession no. GSE113576 and used to create a Seurat object for the analysis. Data were scaled to 10,000 UMI and log10-normalised. Cluster identities were acquired from Supplementary Table 1. accompanying the original publication (<https://science.sciencemag.org/content/362/6416/eaau5324>). Neuronal cluster identities were taken from ‘Neuronal cluster (determined from clustering of inhibitory or excitatory neurons)’ annotations. Data were run through the workflow, once with only POA neurons (removing the 9 suspected extra-POA groups) with upregulated genes capped at 2FC.

**Mouse Brain Atlas (Zeisel et al., 2018)**

Zeisel et al. (2018) created the Mouse Brain Atlas in 2018 – a comprehensive sequencing of the adolescent mouse nervous system. In our previous analyses, data were downloaded from mousebrain.org (<http://mousebrain.org/downloads.html>) and the Level 5 data were downloaded as a one cell per column loom file which also included the cell metadata as column attributes (“l5.all.loom”). Raw data is also available from SRA repository, [SRP135960](https://www.ncbi.nlm.nih.gov/sra/SRP135960)

**Whole Hypothalamus (Chen, Wu, Jiang, and Zhang (2017))**

Chen et al. (2017) sequenced adult (8-10 weeks) whole hypothalamus of female B6D2F1 mice (C57B6 female × DBA2 male). In our previous analyses, processed/normalised data were downloaded as an R object from Gene Expression Omnibus through accession no. [GSE87544](https://www.ncbi.nlm.nih.gov/geo/query/acc.cgi?acc=GSE87544) alongside count data and a metadata file with the cell identities.

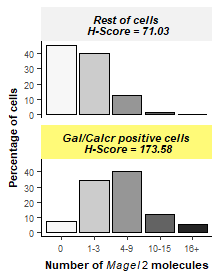

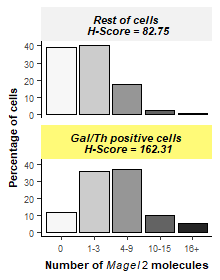

**a**

**b**

**Supplemental Figure S1.** (A) Histogram showing the percentage of cells with particular number of *Magel2* molecules in *Gal/Th* positive cells vs. all other cells. A larger percentage of *Gal/Th* cells expressed 4+ *Magel2* RNA molecules and a smaller percentage of cells expressed zero *Magel2* RNA molecules which contributed to the differences in H-Score. (B) Histogram showing the percentage of cells with particular number of *Magel2* molecules in *Gal/Calcr* positive cells vs. all other cells. A larger percentage of *Gal/Calcr* cells expressed 4+ *Magel2* RNA molecules and a smaller percentage of cells expressed zero *Magel2* RNA molecules which contributed to the differences in H-Score.

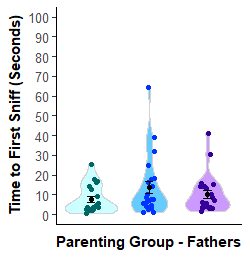

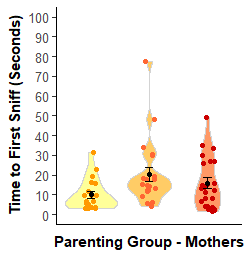

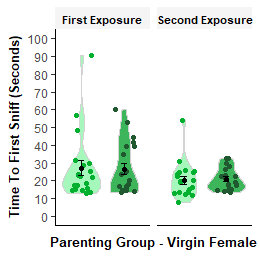

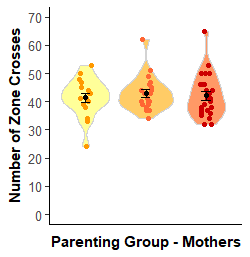

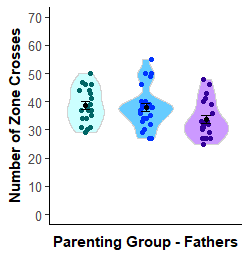

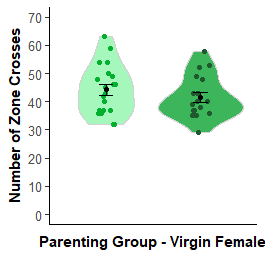

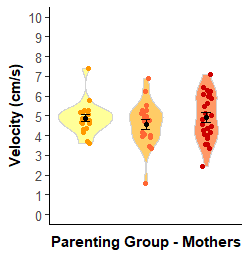

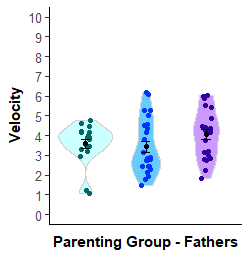

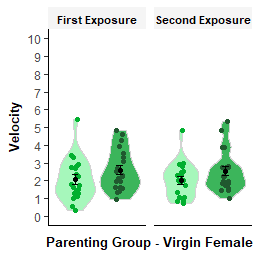

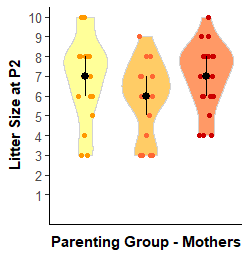

**a**

**b**

**c**

**d**

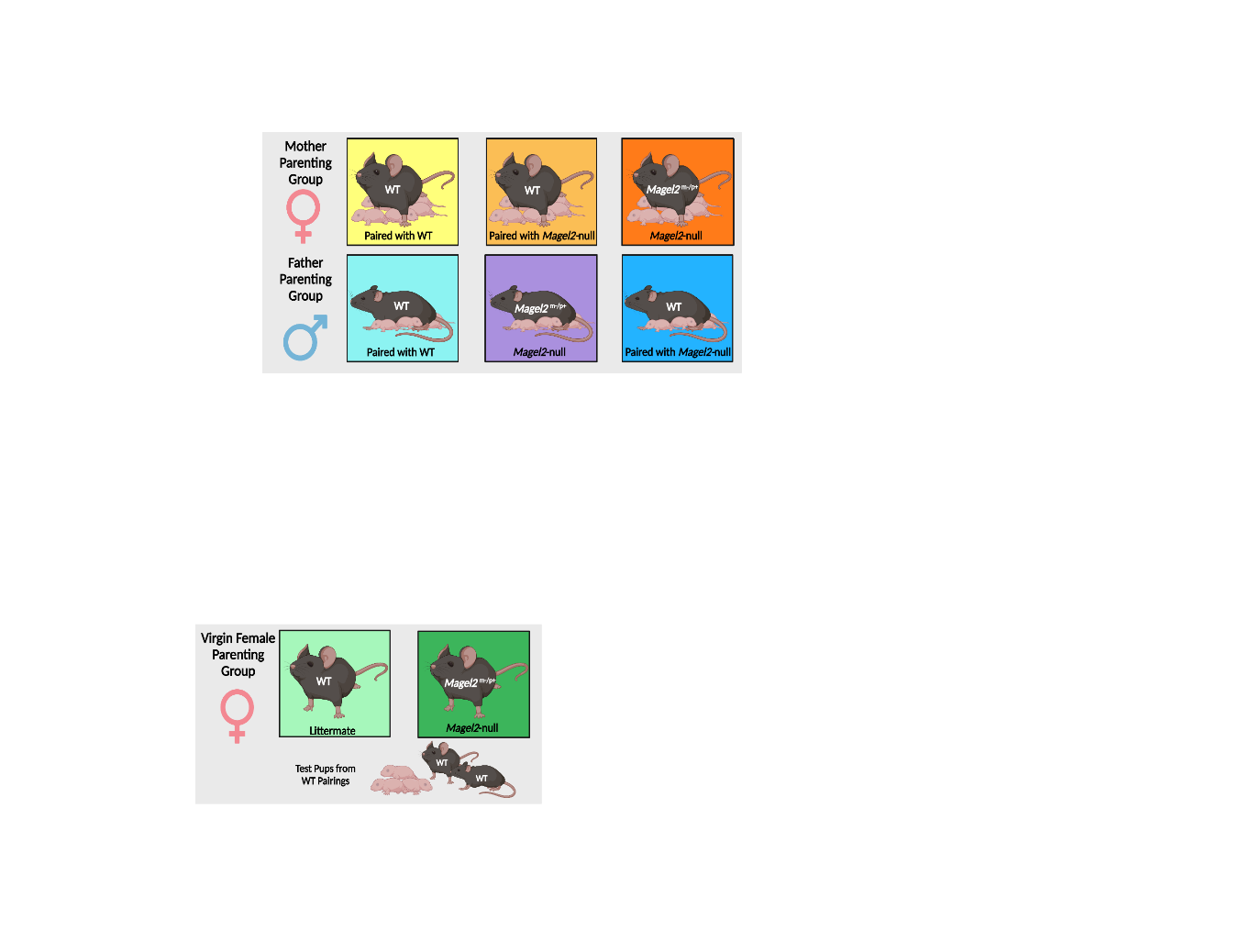

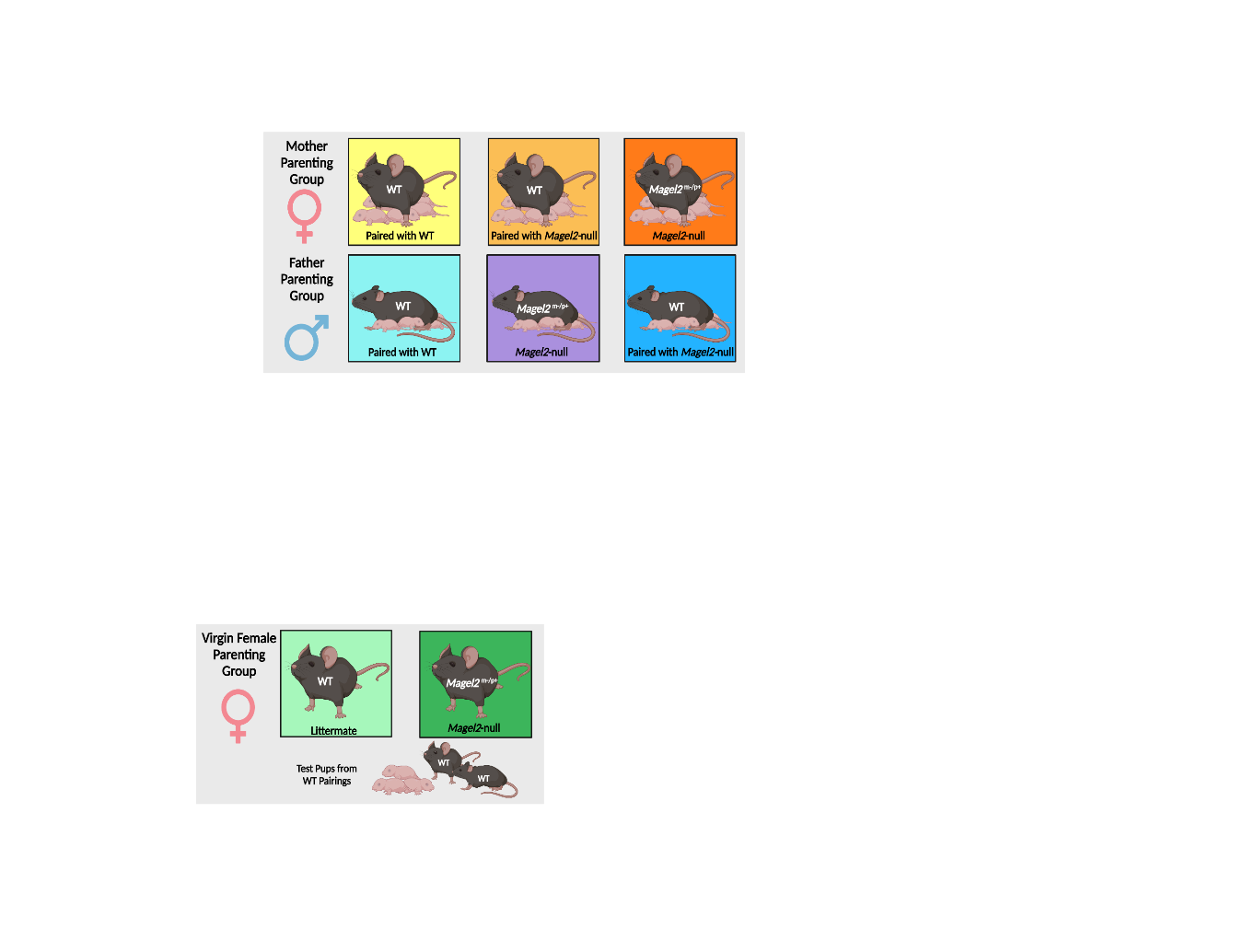

**Supplemental Figure S2. (Previous Page)** Confounders of parenting measured during the retrieval and nest building task and the three chambers task in mothers, fathers and virgin females. (A) Litter size recorded at P2 for the mother/father pairings. (B) Time taken to first sniff a pup in the retrieval and nest building taskfor all three groups of animals. (C) Velocity of each group of animals during the 60-minute retrieval and nest building task measured via automated tracking on Ethovision. (D) The number of chamber crosses in the three-chamber assessment of the three groups of animals measured via automated tracking on Ethovision.

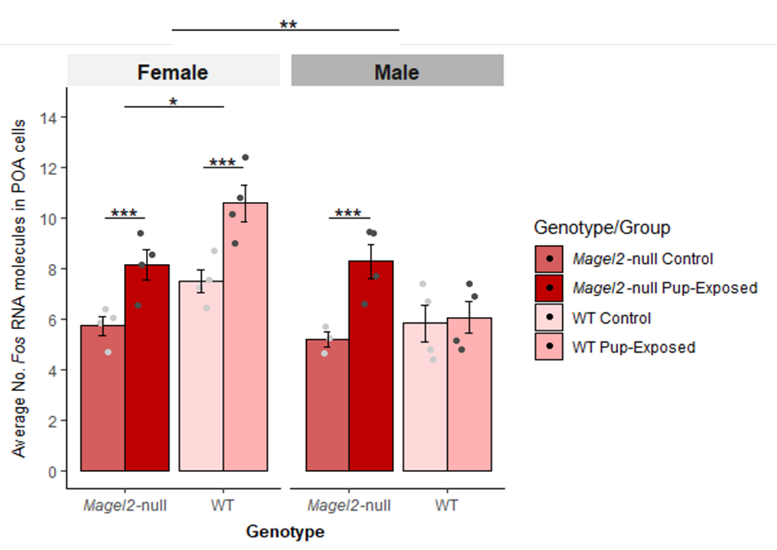

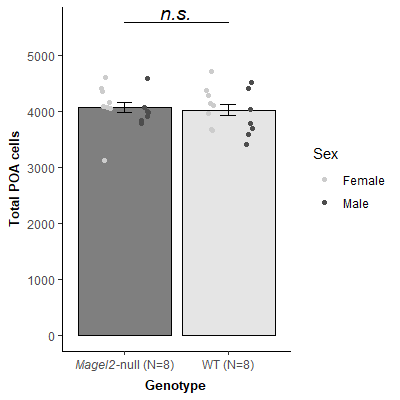

**b**

**a**

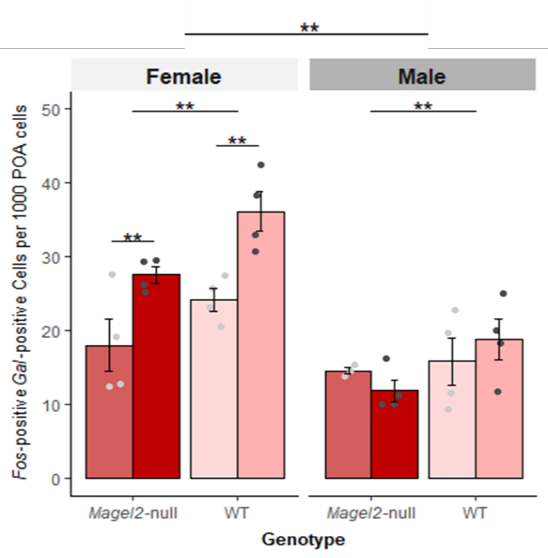

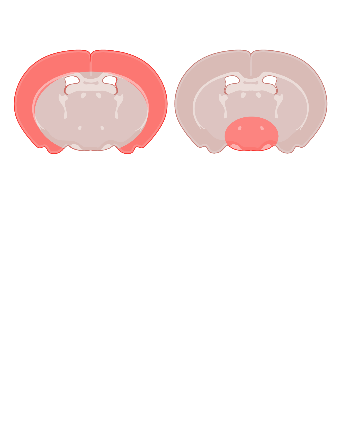

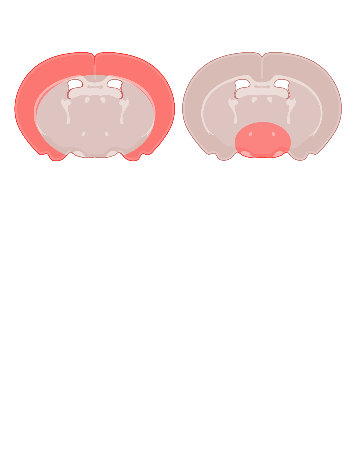

**c**

**d**

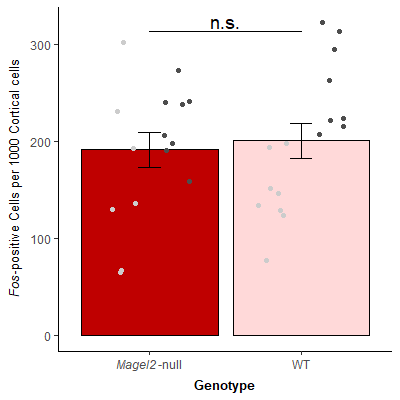

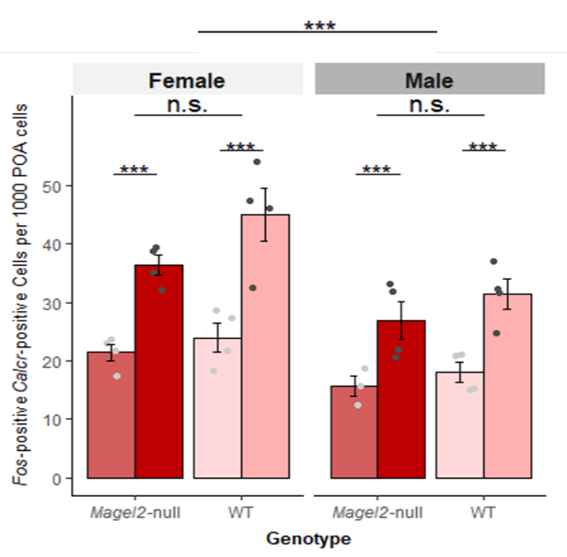

**e**

**f**

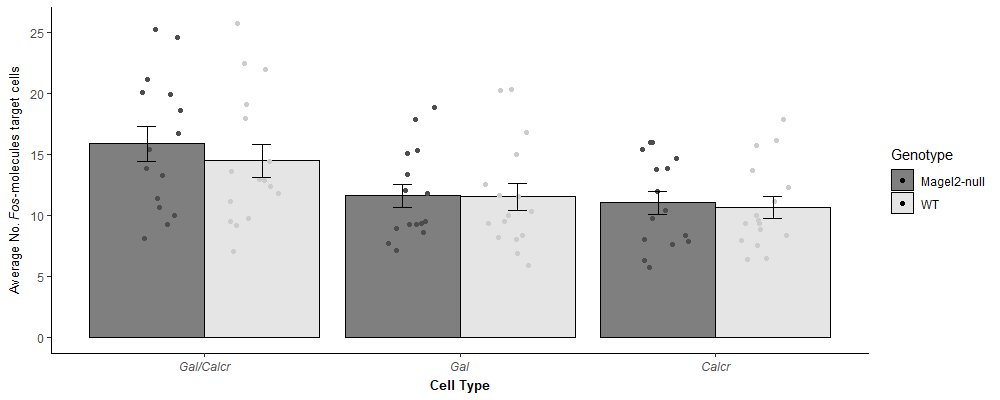

**Supplemental Figure S3.** (A) Number of total POA cells per animal used in this study, split by *Magel2*-null and WT (regardless of condition (B) Average number of *c-Fos* molecules present in POA cells of *Magel2*-null mice and WT mice either exposed to pups or controls. (C) Number of *c-Fos* positive (5+ molecules) cells per 1000 Cortical cells (left) and per 1000 POA cells (right) in *Magel2*-null animals vs. WT animals (regardless of exposure). (D) Number of *c-Fos* positive (5+ molecules) cells also expressing *Gal* (2+ molecules) per 1000 POA cells of *Magel2*-null mice and WT mice either exposed to pups or controls. (E) Number of *c-Fos* positive (5+ molecules) cells also expressing *Calcr* (2+ molecules) per 1000 POA cells of *Magel2*-null mice and WT mice either exposed to pups or controls. (F) Average number of *c-Fos* molecules in POA cell also expressing *Gal/Calcr*, *Gal* and *Calcr*, split by *Magel2*-null and WT (regardless of condition).

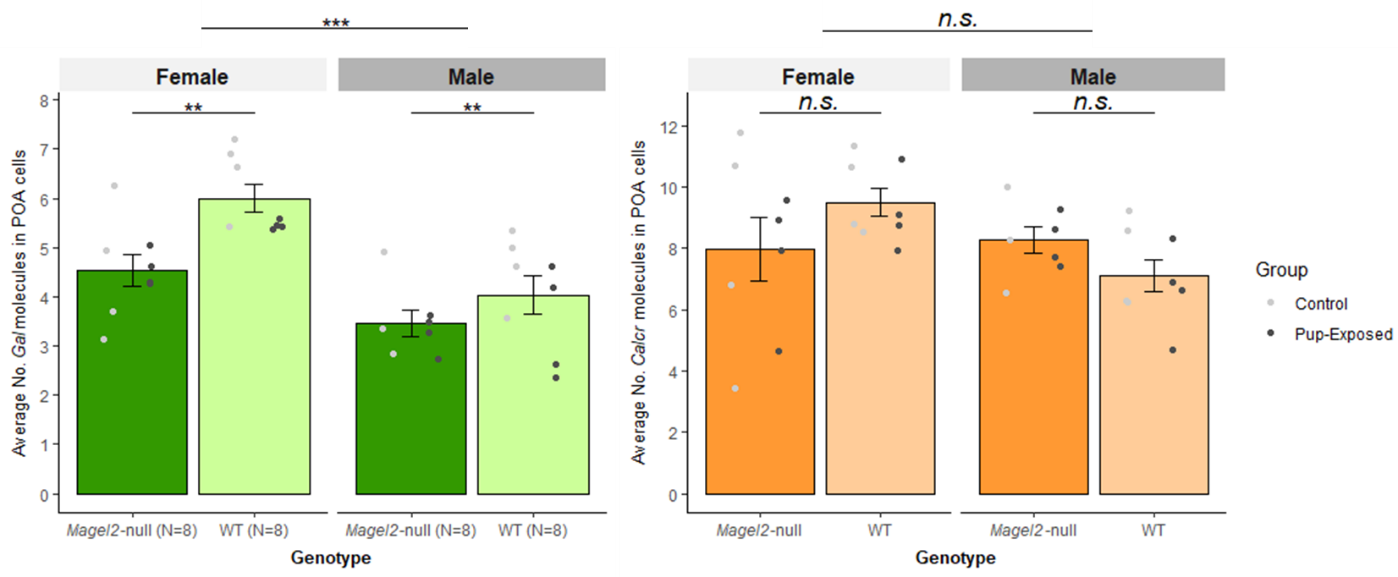

**a**

**b**

**c**

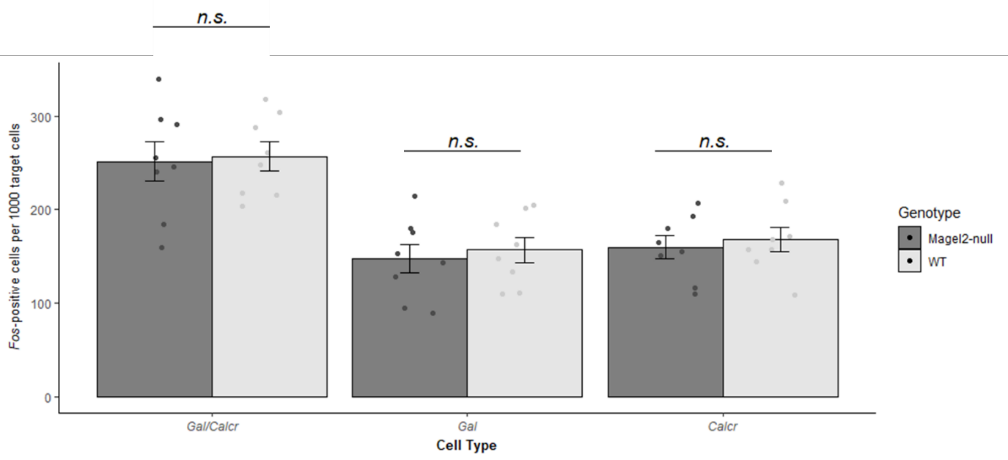

**Supplemental Figure S4.** (A) Average number of *Gal* molecules present in POA cells of *Magel2*-null animals vs. WT animals (regardless of exposure). (B) Average number of *Calcr* molecules present in POA cells of *Magel2*-null animals vs. WT animals (regardless of exposure). (C) Number of (*Gal/Calcr, Gal, Calcr*) cells also registering as *C-Fos* positive (5+ molecules) per 1000 of the respective cell (*Gal/Calcr, Gal, Calcr*). Significant differences seen in Figure 7D & S3D are no longer present here when normalising for *Gal* and *Gal/Calcr* cell number.

| Supplemental Table S1. Specific Hypothalamic Nuclei Level Enrichment Analysis for all imprinted genes in neuronal subpopulation in the Preoptic Area (Moffitt et al., 2018) *Up Reg* – number of upregulated genes with *q* ≤ 0.05 and Log2FC ≥ 1 (total number of genes in the dataset in brackets); *IG* – number of imprinted genes upregulated with *q* ≤ 0.05 and Log2FC ≥ 1 (total number of IGs in the dataset in brackets); *ORA p* – *p* value from over representation analysis on groups with minimum 5% of total IGs; *ORA q* – Bonferroni corrected *p* value from ORA; *Mean FC IG* – mean fold change for upregulated imprinted genes; *Mean FC Rest* – mean fold change for all other upregulated genes; *GSEA p* – *p* value from Gene Set Enrichment Analysis for identity groups with 15+ IGs and Mean FC IG > Mean FC Rest; *GSEA q* – Bonferroni corrected *p* values from GSEA | | | | | | | |
| --- | --- | --- | --- | --- | --- | --- | --- |
| POA Neuron Identity | **Up Reg**  **(16,402)** | **IG**  **(100)** | **ORA *p*** | **ORA *q*** | **Mean FC**  **IG** | **Mean FC**  **Rest** | **GSEA *p*** |
| i16:*Gal/Th* | 825 | 15 | 0.000137 | **0.002594** | 4.89 | 4.16 | 0.416 |
| i35:*Crh/Tac2* | 393 | 9 | 0.000662 | **0.012572** | 2.84 | 5.20 | - |
| i8:*Gal/Amigo2* | 353 | 7 | 0.005834 | 0.110848 | 3.81 | 3.67 | - |
| e13:*Ghrh/C1ql1* | 1088 | 14 | 0.006165 | 0.117129 | 4.28 | 5.79 | - |
| e4:*Trh/Angpt1* | 606 | 9 | 0.011653 | 0.221404 | 3.23 | 5.41 | - |
| e15:*Ucn3/Brs3* | 433 | 7 | 0.016667 | 0.316681 | 13.73 | 5.98 | - |
| e17:*Th/Adcyap1* | 1659 | 17 | 0.02239 | 0.425404 | 4.39 | 5.21 | - |
| i23:*Crh/Nts* | 270 | 5 | 0.024755 | 0.470349 | 2.66 | 4.22 | - |
| e22:*Gal/Ucn3* | 1033 | 11 | 0.049722 | 0.944712 | 3.89 | 10.00 | - |
| i44:*Th/Cxcl14* | 557 | 7 | 0.053803 | 1 | 8.31 | 6.63 | - |
| i37:*Bdnf/Chrm2* | 806 | 9 | 0.057269 | 1 | 5.34 | 5.74 | - |
| i22:*Gal/Pmaip1* | 417 | 5 | 0.111348 | 1 | 4.76 | 4.26 | - |
| e3:*Cartpt/Isl1* | 799 | 8 | 0.11433 | 1 | 8.73 | 4.94 | - |
| e19:*Ghrh/Trh* | 937 | 9 | 0.117593 | 1 | 3.23 | 6.39 | - |
| e2:*Tac1/Fezf1* | 569 | 6 | 0.134146 | 1 | 2.77 | 4.13 | - |
| i24:*Nmu* | 610 | 6 | 0.168524 | 1 | 2.90 | 4.45 | - |
| e10:Glut/*Meis2* | 511 | 5 | 0.201534 | 1 | 3.94 | 5.28 | - |
| e16:*Sst/Cartpt* | 1518 | 12 | 0.212905 | 1 | 3.96 | 5.29 | - |
| e23:*Reln/Etv1* | 1054 | 8 | 0.313293 | 1 | 3.93 | 8.37 | - |
| e1:Glut | 283 | 3 | - | - | 3.39 | 4.22 | - |
| e11:Glut/*Shox2* | 676 | 4 | - | - | 4.44 | 7.37 | - |
| e12:*Nos1/Foxp2* | 482 | 4 | - | - | 3.77 | 5.42 | - |
| e24:*Gal/Rxfp1* | 730 | 4 | - | - | 27.99 | 10.43 | - |
| e5:*Adcyap1/Nkx2-1* | 552 | 2 | - | - | 6.55 | 4.10 | - |
| e6:*Nos1/Trp73* | 368 | 1 | - | - | 17.60 | 5.84 | - |
| e7:*Reln/C1ql1* | 601 | 4 | - | - | 3.40 | 5.18 | - |
| e8:*Cck/Ebf3* | 526 | 1 | - | - | 5.84 | 10.53 | - |
| e9:Glut/*Tcf7l2* | 459 | 1 | - | - | 4.58 | 5.86 | - |
| h1:GABA/*Slc17a6* | 563 | 3 | - | - | 3.30 | 5.99 | - |
| h2:*Nts/Slc17a8* | 237 | 1 | - | - | 2.76 | 4.37 | - |
| i1:GABA | 74 | 0 | - | - | 0.00 | 3.38 | - |
| i10:*Tac1/Nts* | 662 | 3 | - | - | 4.73 | 3.48 | - |
| i11:GABA | 172 | 1 | - | - | 5.43 | 3.69 | - |
| i12:GABA | 177 | 2 | - | - | 6.02 | 4.01 | - |
| i13:GABA | 322 | 2 | - | - | 4.93 | 3.64 | - |
| i15:GABA | 412 | 2 | - | - | 3.26 | 4.97 | - |
| i17:*Th/Nos1* | 290 | 4 | - | - | 4.71 | 4.41 | - |
| i18:*Gal/Tac2* | 516 | 4 | - | - | 2.51 | 3.75 | - |
| i2:*Tac1/Pdyn* | 620 | 4 | - | - | 4.65 | 3.63 | - |
| i20:*Gal/Moxd1* | 444 | 4 | - | - | 3.47 | 5.26 | - |
| i21:*Sst/Pou3f3* | 602 | 4 | - | - | 4.97 | 4.21 | - |
| i25:*Npy/Etv1* | 396 | 4 | - | - | 4.19 | 6.07 | - |
| i26:*Tac1/Prok2* | 110 | 1 | - | - | 7.04 | 5.15 | - |
| i29:GABA*/Igsf1* | 418 | 2 | - | - | 2.52 | 5.21 | - |
| i3:*Penk/Nts* | 591 | 3 | - | - | 2.95 | 3.99 | - |
| i32:*Sst/Npy* | 350 | 2 | - | - | 3.14 | 7.38 | - |
| i38: *Kiss1/Th* | 636 | 3 | - | - | 6.45 | 13.13 | - |
| i39:GABA | 331 | 2 | - | - | 2.74 | 7.13 | - |
| i4:GABA/*Mylk* | 232 | 2 | - | - | 2.95 | 4.53 | - |
| i40:*Sst/Reln* | 270 | 3 | - | - | 7.75 | 7.55 | - |
| i41:*Npy/Penk* | 376 | 4 | - | - | 7.15 | 14.72 | - |
| i42:*Pthlh* | 375 | 3 | - | - | 5.75 | 8.11 | - |
| i43*:Chat* | 1174 | 3 | - | - | 11.91 | 13.93 | - |
| i45:*Bdnf/Pmaip1* | 182 | 1 | - | - | 23.06 | 12.12 | - |
| i5:GABA/*Pou3f3* | 525 | 1 | - | - | 3.23 | 3.93 | - |
| i7:GABA | 261 | 2 | - | - | 4.52 | 4.42 | - |
| i9:GABA | 284 | 1 | - | - | 2.09 | 4.12 | - |

| Supplemental Table S2A. Top 5 neuronal subpopulations showing imprinted gene over-representation in the MBA dataset (Zeisel et al., 2018). *Neuron Description –* more specific classification of neuronal subpopulations as described by Zeisel et al. (2018); All column descriptions can be found in the legend of Supplemental Table S1 | | | | | | | | | |
| --- | --- | --- | --- | --- | --- | --- | --- | --- | --- |
| Neuron Identity | **Neuron**  **Description** | **Up Reg**  **(18,335)** | **IG**  **(105)** | **ORA *p*** | **ORA *q*** | **Mean FC**  **IG** | **Mean FC**  **Rest** | **GSEA**  ***p*** | **GSEA**  ***q*** |
| HBSER5 | Serotonergic neurons, hindbrain | 3721 | 53 | 8.29E-12 | **1.19E-09** | 13.76 | 5.98 | 0.002 | **0.042** |
| DEINH5 | Peptidergic neurons, hypothalamus | 611 | 19 | 1.94E-09 | **2.79E-07** | 7.47 | 5.48 | 0.0639 | 1 |
| TEINH3 | Inhibitory neurons, telencephalon | 885 | 22 | 5.50E-09 | **7.93E-07** | 8.60 | 6.38 | 0.2076 | 1 |
| MEINH13 | Inhibitory neurons, midbrain | 1229 | 25 | 2.35E-08 | **3.38E-06** | 9.85 | 7.30 | 0.1213 | 1 |
| HBSER4 | Serotonergic neurons, hindbrain | 3510 | 45 | 2.83E-08 | **4.08E-06** | 27.93 | 6.29 | 5.00E-04 | **0.0105** |

| Supplemental Table S2B. Whole Hypothalamus Level Enrichment Analysis for all imprinted genes in neuronal subpopulations in the Chen et al. (2017) dataset. All column descriptions can be found in the legend of Supplemental Table S1. Marker Genes – Genes identified by Chen et al. (2017) with distinct expression in these neuronal subtypes for the purposes of identification. | | | | | | | |
| --- | --- | --- | --- | --- | --- | --- | --- |
| Hypothalamic Neuron Identity | **Marker**  **Genes** | **Up Reg**  **(12,238)** | **IG**  **(80)** | **ORA *p*** | **ORA *q*** | **Mean FC**  **IG** | **Mean FC**  **Rest** |
| GABA17 | *Slc6a3* | 825 | 18 | 4.57E-06 | **1.19E-04** | 2.73 | 3.30 |
| GABA8 | *Vipr2* | 480 | 11 | 0.0003 | **0.0071** | 1.96 | 2.73 |
| GABA13 | *Slc18a2, Gal* | 569 | 12 | 0.0003 | **0.0079** | 1.89 | 2.02 |
| GABA15 | *Agrp* | 766 | 13 | 0.0013 | **0.0339** | 2.40 | 2.79 |
| GABA16 | *Cox6a2* | 258 | 6 | 0.0068 | 0.1762 | 1.68 | 2.70 |
| GABA14 | *Cbln4* | 271 | 6 | 0.0085 | 0.2220 | 1.68 | 2.54 |
| GABA2 | *Npas1* | 278 | 6 | 0.0096 | 0.2499 | 2.29 | 4.22 |
| GABA18 | *Lhx1, Klhl1* | 200 | 5 | 0.0099 | 0.2576 | 1.55 | 3.06 |
| GABA9 | *Vip* | 219 | 5 | 0.0142 | 0.3703 | 1.39 | 3.77 |
| GABA10 | *Prok2* | 605 | 9 | 0.0169 | 0.4391 | 3.03 | 4.28 |
| Glu9 | *Gng8, Samd3* | 345 | 6 | 0.0252 | 0.6556 | 1.34 | 3.66 |
| Glu1 | *Crh* | 182 | 4 | 0.0312 | 0.8120 | 1.68 | 4.38 |
| GABA11 | *Ghrh* | 392 | 6 | 0.0430 | 1 | 6.50 | 2.91 |
| GABA5 | *Lhx8* | 569 | 7 | 0.0780 | 1 | 1.28 | 3.09 |
| Glu14 | *Avp, Sim1* | 945 | 10 | 0.0875 | 1 | 2.07 | 3.30 |
| Glu15 | *Sst, Prdm8* | 650 | 7 | 0.1319 | 1 | 1.69 | 3.79 |
| Glu12 | *Vgll2* | 339 | 4 | 0.1811 | 1 | 1.70 | 3.88 |
| Glu13 | *Pomc* | 471 | 5 | 0.1945 | 1 | 3.36 | 4.04 |
| GABA12 | *Crabp1* | 354 | 4 | 0.2010 | 1 | 1.63 | 2.61 |
| Glu6 | *Tac1* | 607 | 6 | 0.2052 | 1 | 2.39 | 3.58 |
| Glu7 | *Fezf1, Lbhd2* | 609 | 6 | 0.2072 | 1 | 1.44 | 2.13 |
| Glu11 | *Kiss1* | 468 | 4 | 0.3663 | 1 | 7.78 | 3.68 |
| GABA3 | *Bcl11b* | 484 | 4 | 0.3902 | 1 | 1.41 | 3.36 |
| Glu8 | *Lbhd2, Cartpt* | 534 | 4 | 0.4641 | 1 | 3.34 | 2.97 |
| Glu4 | *Shox2* | 1690 | 8 | 0.8792 | 1 | 2.25 | 3.09 |
| Glu5 | *Foxb1* | 984 | 4 | 0.8942 | 1 | 2.20 | 3.69 |
| GABA6 | *Pax6* | 228 | 3 | - | - | 1.03 | 3.86 |
| GABA7 | *Trh* | 194 | 3 | - | - | 3.87 | 3.46 |
| Glu3 | *Fezf2, Samd3* | 203 | 3 | - | - | 1.07 | 5.35 |
| GABA1 | *Pvalb* | 738 | 2 | - | - | 3.81 | 4.57 |
| GABA4 | *Gm13498* | 254 | 2 | - | - | 3.97 | 4.43 |
| Glu10 | *Trh* | 228 | 2 | - | - | 1.17 | 4.88 |
| Hista | *Hdc* | 585 | 2 | - | - | 3.98 | 5.14 |
| Glu2 | *Sln* | 376 | 0 | - | - | 0.00 | 4.38 |

**Supplemental Table S3.** Marker Genes of the Galanin Neuron Subtypes in Zeisel et al. (2018) and Chen et al. (2017) datasets.

(Supplemental_Table_S3_Marker_Genes.xlsx)

| **Supplemental Table S4A.** Outcome of statistical tests for *Gal/Th* and *Gal/Calcr* *Magel2* expression comparisons. For the *Gal/Th* (top) and *Gal/Calcr* (bottom) experiments, below are the results of comparing the number of *Magel2* molecules present in these cells to the rest of all POA cells, the other *Gal* positive POA cells as well as the *Th/Calcr* positive cells. Finally, a comparison was made when removing all Magel2 non-expressing cells from the analysis and so assessing whether *Magel2* was more strongly expressed in target cells vs. other *Magel2* expressing cells. | | | | | | |
| --- | --- | --- | --- | --- | --- | --- |
| **Group (N)** | **No. of Cells** | **% Cells *Magel2* Positive** | **Avg. No. *Magel2* molecules** | **Statistical Test** | **Test statistic vs. target group** | ***p* value vs. target group** |
| ***Gal/Th* POA (N=2) compared to ...** | 2579 | 88.29% | 5.28 |  |  |  |
| **Rest of All POA Cells (N=2)** | 66,981 | 61.26% | 2.11 | Mann Whitney U | W = 47192483 | < 2.2x10-16 |
| **Rest of POA *Gal* (N=2)** | 2,511 | 82.99% | 4.11 | Bonferroni Corrected Dunn Test | Z = 6.87 | 3.85x10-11 |
| **Rest of POA *Th* (N=2)** | 12,912 | 77.17% | 3.22 | Bonferroni Corrected Dunn Test | Z = 19.91 | 2.14x10-87 |
| **Rest of POA Cells (No *Gal*/No *Th*) (N=2)** | 51,558 | 56.22% | 1.74 | Bonferroni Corrected Dunn Test | Z = 46.5 | 0.00 |
| ***Gal/Th* *Magel2* +ve POA (N=2) compared to …** | 2,277 | 100.00% | 5.99 |  |  |  |
| **Rest of All POA Cells *Magel2* +ve (N=2)** | 41,033 | 100.00% | 3.45 | Mann Whitney U | W = 30882369 | < 2.2x10-16 |
| ***Gal/Calcr* POA (N=4) compared to …** | 3846 | 92.23% | 5.81 |  |  |  |
| **Rest of All POA Cells (N=4)** | 152,323 | 54.48% | 1.59 | Mann Whitney U | W = 110992285 | < 2.2x10-16 |
| **Rest of POA *Gal* (N=4)** | 23,512 | 76.72% | 3.06 | Bonferroni Corrected Dunn Test | Z = 32.64 | 7.34x10-233 |
| **Rest of POA *Calcr* (N=4)** | 4,818 | 84.81% | 3.30 | Bonferroni Corrected Dunn Test | Z = 19.08 | 3.46x10-81 |
| **Rest of POA *Cells* (No *Gal*/No *Calcr*) (N=4)** | 123,993 | 49.09% | 1.24 | Bonferroni Corrected Dunn Test | Z = 77.54 | 0.00 |
| ***Gal/Calcr* Magel2 +ve POA (N=4) compared to …** | 3,547 | 100% | 6.30 |  |  |  |
| **Rest of All POA Cells *Magel2* +ve (N=4)** | 82,990 | 100% | 2.91 | Mann Whitney U | W = 75812992 | < 2.2x10-16 |

| **Supplemental Table S4B.** Proportion of cells with varying numbers of Magel2 reads and the corresponding H-scores. For the experiments in Chapter 4, the proportion of different cell types in the POA/PVN which have a binned number of Magel2 RNA molecules. H-scores are calculated from weighted calculation of proportions | | | | | | | |
| --- | --- | --- | --- | --- | --- | --- | --- |
| ***Cell Type*** | **0 *Magel2* reads** | ***1-3 Magel2* reads** | **4-9 *Magel2* reads** | **10-15 *Magel2* reads** | **16+ *Magel2* reads** | **H-Score** | **H-score - only expressors** |
| **POA - *Gal/Calcr*** | | | | | | | |
| ***Gal/Calcr*** | 0.08 | 0.37 | 0.43 | 0.13 | 0.06 | 173.583 | 188.215 |
| ***Gal*** | 0.30 | 0.59 | 0.33 | 0.06 | 0.02 | 115.379 | 150.385 |
| ***Calcr*** | 0.18 | 0.60 | 0.34 | 0.05 | 0.01 | 125.633 | 148.140 |
| **Rest** | 0.84 | 0.74 | 0.23 | 0.03 | 0.01 | 71.029 | 130.370 |
| **POA - *Gal/Th*** | | | | | | | |
| ***Gal/Th*** | 0.13 | 0.40 | 0.42 | 0.12 | 0.06 | 162.311 | 183.838 |
| ***Gal*** | 0.20 | 0.47 | 0.40 | 0.10 | 0.03 | 85.941 | 168.762 |
| ***Th*** | 0.30 | 0.58 | 0.34 | 0.06 | 0.02 | 140.064 | 152.579 |
| **Rest** | 0.63 | 0.66 | 0.29 | 0.04 | 0.01 | 117.743 | 140.287 |

| Supplemental Table S5. Statistical results from mother and father analyses carried out without including parents with test litters containing mutant pups. Statistical tests used are the same as reported in the main text P values in red indicate disparities with the original statistical analysis. | | | | | |
| --- | --- | --- | --- | --- | --- |
| Mothers (N=11) | | | **Pairwise p** | | |
| Measure | **Test Stat** | **p** | **WT(WT) vs. WT(HET)** | **WT(WT) vs. HET** | **WT(HET) vs. HET** |
| Task Fin | 20.72 | **2.59x10^-7^** | 1 | **6.7x10^-7^** | **0.00021** |
| Task Status | 15.727 | **0.00038** | 1 | **0.001** | **0.01** |
| No. Pups Ret | NA | NA | NA | NA | NA |
| First Ret | 1.34 | 0.271 | NA | NA | NA |
| Last Ret | 2.804 | 0.069 | NA | NA | NA |
| Level 3 Nest | 20.72 | **2.59x10^-7^** | 1 | **6.7x10^-7^** | **0.00021** |
| Nest Quality | 17.363 | **0.00017** | 1 | **0.0003** | **0.015** |
| PDB Ret | 6.862 | **0.002** | 1 | **0.0043** | **0.035** |
| PDB Fin | 19.26 | **5.88x10^-7^** | 1 | **1.8x10^-6^** | **0.00024** |
| Fathers (N=11) | | | **Pairwise p** | | |
| Measure | **Test Stat** | **p** | **WT(WT) vs. WT(HET)** | **WT(WT) vs. HET** | **WT(HET) vs. HET** |
| Task Fin | 14.771 | **0.0006** | 0.676 | **0.0004** | **0.0093** |
| Task Status | 22.391 | **1.374 x 10^-5^** | 0.88 | **1.4x10^-5^** | **0.0003** |
| No. Pups Ret | 18.315 | **0.0001** | 1 | **0.0002** | **0.00038** |
| First Ret | 6.53 | **0.003** | 1 | **0.0026** | **0.0159** |
| Last Ret | 15.833 | **0.0004** | 1 | **0.0005** | **0.001** |
| Level 3 Nest | 16.512 | **0.0003** | 0.19 | **0.0001** | **0.02** |
| Nest Quality | 11.327 | **0.003** | **0.044** | **0.004** | 0.61 |
| PDB Ret | 4.597 | **0.0145** | 1 | **0.014** | **0.052** |
| PDB Fin | 6.29 | **0.0036** | 0.466 | **0.0025** | **0.047** |

**Supplemental Table S6.** Gal/Calcr/Fos RNAscope® ANOVA Outputs. Outputs of all 3-way and 2-way ANOVA’s performed from the Gal/Calcr/Fos RNAscope® experiments

(‘Supplemental_Table_S6_GalCalcrFos_ANOVA_Output’)

**Supplemental Table S7.** Imprinted Gene List. Custom List of Imprinted Genes used as the gene set in this study.

(‘Supplemental_Table_S7_Imprinted_Gene_List.xlsx')

| Supplemental Table S8. Intra class correlation coefficient (ICC) Kappa and *p* values as measures of the primary and second scorer inter-rater reliability. Statistics were calculated from the 80/210 videos that were second scored | | |
| --- | --- | --- |
| Metric from Retrieval/Nest Building Assessment | **ICC Kappa (0-1)** | **ICC *p* value** |
| *Time to Retrieve Pup 1* | 0.915 | 1.89x10^-33^ |
| *Time to Retrieve Pup 2* | 0.936 | 3.05x10^-38^ |
| *Time to Retrieve Pup 3* | 0.958 | 3.05x10^-45^ |
| *Number of Pups Retrieved* | 0.935 | 6.12x10^-38^ |
| *Time to Build a Level 3 Nest* | 0.894 | 7.44x10^-30^ |
| *Final Nest Quality Rating* | 0.778 | 4.61x10^-18^ |
| *Time Until Task Finished* | 0.872 | 9.33x10^-27^ |
| *PDB Until Retrieval* | 0.853 | 2.36x10^-24^ |
| *PDB Until Task Finished* | 0.928 | 3.46x10^-36^ |

| **Supplemental Table S9.** Settings used for each gene/probe combination when acquiring images used Zeiss AxioScan Z1. Gene/probe combinations are listed with the catalogue code the probe was registered under in the ACD Biotechne catalogue and the specific experiment the probe was used in within this thesis. For each probe combination, the intensity percentage and light duration (milliseconds (ms)) are recorded. | | | | |
| --- | --- | --- | --- | --- |
| **Experiment** | **Gene / Probe** | **Probe Catalogue code** | **HXP**  **120 V Intensity** | **HXP**  **120 V Duration** |
| **TSA Vivid**^TM^ **520 (Green, Fluorescein)** | | | | |
| *Calcr/Gal/Magel2* | *Calcr* | ACD 494071,Mm-Calcr | 50% | 1500ms |
| *Gal/Calcr/Fos & Gal/Th/Magel2* | *Gal* | ACD 400961, Mm-*Gal* | 50% | 400ms |
| *Oxt/Avp/Magel2* | *Oxt* | ACD 493171 Mm-*Oxt* | 50% | 100ms |
| **TSA Vivid**^TM^ **570 (Orange, Cy3)** | | | | |
| *Calcr/Gal/Magel2* | *Gal* | ACD 400961-C2, Mm-*Gal*-C2 | 100% | 15ms |
| *Gal/Th/Magel2* | *Th* | ACD 317621-C2, Mm-*Th*-C2 | 100% | 25ms |
| *Gal/Calcr/Fos* | *Calcr* | ACD 494071-C2,Mm-Calcr-C2 | 100% | 10ms |
| *Oxt/Avp/Magel2* | *Avp* | ACD 401391-C2, Mm-*Avp*-C2 | 100% | 2.5ms |
| **TSA Vivid**^TM^ **650 (Red, Cy5.5)** | | | | |
| *Calcr/Gal/Magel2, Gal/Th/Magel2, Oxt/Avp/Magel2* | *Magel2* | ACD 502971-C3, Mm-*Magel2*-C3 | 100% | 150ms |
| *Gal/Calcr/Fos* | *Fos* | ACD 316921-C3, Mm-*Fos*-C3 | 100% | 200ms |
